# RNA-directed DNA methylation controls seed development and heat stress memory in barley

**DOI:** 10.64898/2026.07.30.741739

**Authors:** Auwalu Abdu, Henrik Mihály Szaker, András Kis, Dávid Polgári, Ágnes Dalmadi, Marianna Rakszegi, Tibor Csorba, Zoltán Havelda

## Abstract

Plant specific RNA-directed DNA methylation (RdDM) mediates the DNA methylation of specific DNA sequences directed by 24-nucleotide long (nt) small interfering (si)RNAs. In crop plants we have limited information about the biological roles of the RdDM pathway including barley (*Hordeum vulgare* L). Here, we show that knockout of barley *NRPD/E2A* gene by genome editing, bringing about the drastic inhibition of RdDM pathway, results in the early arrest of developing caryopses rendering the mutant plants sterile. The knockout of the downstream component *RDR2* gene, responsible for generating double stranded precursor RNAs for the production of 24-nt siRNAs, was associated with the loss of the majority of these siRNAs and also typically induced the early arrest of caryopsis development. However, the *rdr2* mutants were able to produce a limited number of seeds exhibiting smaller size, endosperm filling anomalies and inhibited germination, which phenomenon was predominantly inherited maternally. Genome-wide analyses of gene expression revealed drastic up-and down-regulations in *rdr2* mutants compared to the wild type. We did not find direct correlation between the changes of gene expression and the localization of 24-nt siRNA producing clusters indicating the indirect action of RdDM in these regulatory events. We also demonstrate that *rdr2* mutant shows reduced heat stress memory capacity rendering the mutant plants more vulnerable to high temperature. Altogether, our data show the RdDM pathway is a major regulatory contributor to generative development and heat stress responses in barley.

## Introduction

RNA silencing (or RNA interference, RNAi) is a highly conserved gene regulatory mechanism acting in eukaryotic organisms. RNA silencing functions in a sequence specific manner via the action of small regulatory RNAs (sRNAs) of 20-25-nucleotide (nt) length. sRNAs control the level or translation capacity of their target RNAs which are involved in a wide range of biological processes. sRNAs can be categorized into two main groups, the microRNAs (miRNAs) and the small interfering RNAs (siRNAs), the latter having several sRNA subclasses including heterochromatic siRNA (het-siRNAs), or repeat-associated siRNAs (ra-siRNAs), in case of having derived from repetitive elements (Vaucheret and Voinnet, 2024). MiRNAs (21-nt long) typically control euchromatic gene expression while in plants, het-siRNAs of 24-nt length, are typically thought to orchestrate peri/centromeric and (sub)telomeric heterochromatin structures (Vrbsky et al., 2010), preventing the disruptive movement of mobile elements both within heterochromatic blocks and interspersed euchromatic regions through RNA-directed DNA methylation (RdDM) (Erdmann and Picard, 2020). In plants, similarly to other eukaryotes, DNA methylation acts as an epigenetic mark, playing a pivotal role in the silencing of transposable elements (TEs) and hence maintaining genome stability, as well as the regulation of gene expression, development, and environmental responses (Zhang et al., 2018). However, DNA methylation in plants differs from the mammalian systems regarding the sequence context, genomic location, and inheritance stability. Animal genomes are predominantly methylated at the symmetric GC context, while in plants DNA methylation occurs in three sequence contexts, CG, CHG and CHH (H meaning A, T, or C) (Xie et al., 2025). Crucially, while animals lack the RdDM pathway and instead utilize piRNAs for small RNA-guided TE silencing, plants rely heavily on 24-nt het-siRNAs and plant-specific RNA polymerases to target *de novo* methylation. The regulation of this machinery consists of three steps (Law and Jacobsen, 2010), the *de novo* methylation which establishes the initial pattern, the maintenance of existing methylation and the active DNA demethylation.

The *de novo* establishment of DNA methylation is mediated by RdDM pathway specific for plants (Matzke et al., 2015; Zhang et al., 2022). The canonical RdDM pathway is primed by the action of plant-specific DNA-DEPENDENT RNA POLYMERASE IV (POL IV). In the next step RNA-DEPENDENT RNA POLYMERASE 2 (RDR2) produces a complementary RNA strand of Pol IV transcripts generating double stranded (ds) RNAs (Du, 2024). The DICER-LIKE 3 (DCL3) processes the dsRNAs into 24-nt long siRNAs. One strand of the siRNAs is then loaded into the effector proteins, ARGONAUTE 4 (AGO4) or its paralogs AGO6 or AGO9. AGO complexes loaded with one strand of siRNA are recruited by Watson-Crick base pairing to long non-coding scaffold RNA produced by a second plant specific DNA-DEPENDENT RNA POLYMERASE V (POL V). In *Arabidopsis* the recruited AGO4 complex triggers cytosine methylation through DOMAINS REARRANGED METHYLTRANSFERASE 2 (DRM2) and to a lesser extent its homolog DRM1 in all sequence contexts (Chow and Mosher, 2023). The establishment of methylation facilitates the recruitment of POL IV for additional production of siRNAs creating a stable positive feedback loop. This RdDM feedback loop is important in the reinforcement of non-symmetric CHH DNA methylation which, unlike symmetric contexts, cannot be copied semi-conservatively during DNA replication (Law et al., 2013).

RdDM activities are mainly focused on transposons inducing their transcriptional silencing at the boundaries of heterochromatin and euchromatin (Li et al., 2015; Böhmdorfer et al., 2016). RdDM mediated methylation of transposon sequences can interfere with the expression of protein coding genes in the vicinity (Hollister and Gaut, 2009). In line with the abundant presence of 24-nt siRNAs in reproductive tissues (Mosher et al., 2009)RdDM is mainly required for reproductive development, particularly in reinforcing TE silencing in the germline and regulating genomic imprinting. In addition, RdDM can act via alternative non-canonical pathways engaging distinct mechanisms to modify chromatin (Cuerda-Gil and Slotkin, 2016).

The biological roles of RdDM have been characterized only in a limited number of plant species. In *Arabidopsis* mutations in the RdDM pathway induce only moderate phenotypic alterations such as subtle changes in the seed size and flowering time (Gallego-Bartolome et al., 2019; Kirkbride et al., 2019; Grover et al., 2020). However, in other crops like rice and maize, loss of RdDM components leads to severe developmental defects and sterility, highlighting species-specific dependencies (Moritoh et al., 2012; Zheng et al., 2021; Wang et al., 2022). Furthermore, environmental stress responses, including heat stress (Popova et al., 2013), osmotic stress (Wibowo et al., 2016), phosphate starvation(Secco et al., 2015), or pathogen infection (Dowen et al., 2012)induced transcriptional variations were also connected to the RdDM pathway.

In plants having larger genomes with a high proportion of heterochromatic DNA, the roles of RdDM can be more obvious. The RDR2 mutation in maize modifies the expression pattern of thousands of genes, transposons, and causes significant changes in shoot apical meristem morphology (Jia et al., 2009). The knock-out of an essential component of maize RdDM pathway, KOW DOMAIN-CONTAINING TRANSCRIPTION FACTOR1 (KTF1), affected salt tolerance (Wang et al., 2025). POL IV mutation in *Brassica rapa* induces abortion of seeds after fertilization (Grover et al., 2018). The similar mutation in *Capsella rubella* brings about the post-meiotic arrest of pollen development (Wang et al., 2020). In *Capsella grandiflora,* an obligate out-crosser, the loss of RdDM pathway induced seed abortions, while other self-compatible members of the family showed no defects due to the mutation indicating the importance of mating system in RdDM mutant background (Dew-Budd et al., 2024). In tomato POL IV mutation affects proper embryo development and the improperly maturated embryos are not able to resist post-harvest desiccation (Feng et al., 2025). In rice, the RdDM activity controls economically important traits such as endosperm development, plant height, tiller number and immunity responses (Xu et al., 2024; Pal et al., 2025; Zhang et al., 2026). Furthermore, the RdDM pathway is able to prevent the premature activation of genes associated with immune response in soybean (*Glycine max*) (Wang et al., 2026; Zhang et al., 2026).

Numerous monocot species have agronomic importance and their grain production contributes enormously to global food supply and animal nutrition. To increase their productivity and stress tolerance, the fundamental understanding of development and stress resilience is of utmost importance, since global warming is an imminent and critical threat to global crop production. Barley (*Hordeum vulgare*) is an important monocot crop plant occupying the fourth place in the ranking of production level following wheat, maize, and rice. Worldwide barley grain production reached above 150 M tonnes in 2022/2023 vegetation period (http://faostat3.fao.org). Barley grain is used for animal feeding, production of beverages and human nutrition. Barley, as a self-pollinating diploid plant became also the research model of monocot crops (Harwood, 2019). Despite the extensive research efforts, the biological roles of the RdDM pathway still remains unknown in this economically important crop plant.

Here we show that the RdDM pathway is a pivotal component of barley reproductive development and heat stress resilience, and consequently controlling economically important traits. Uncovering the RdDM mediated regulation of economically important traits opens new avenues for epigenetic breeding strategies. Furthermore, since barley serves as the premier diploid model for hexaploid wheat, our findings elucidating the biological roles of RdDM will also provide valuable insights for wheat breeding programs.

## Results

### The barley *nrpd/e2a* mutants show aborted seed development

To reveal the biological roles of RdDM in barley we have created a series of mutant barley lines disabled in various functions of the RdDM pathway. First, we produced mutants exhibiting a complete loss of RdDM activities. To achieve this goal, we aimed to identify the barley *NRPD/E2* genes encoding the common subunit of POL IV and POL V complexes. Introducing knock-out mutations into these genes is expected to abolish completely the canonical RdDM pathway due to the disruption of both POL IV-dependent siRNA biogenesis and POL V-dependent scaffold RNA transcription (Xie et al., 2024). To identify the barley *NRPD/E2*, first Arabidopsis NRPD/E2 protein sequence was retrieved from the UniProt database and used to run a blast on Ensembl Plants protein database against the barley ‘Golden Promise’. We identified two paralogs of *NRPD/E2* gene in barley, which we designated *HvNRPD/E2A* and *HvNRPD/E2B* (Supplementary Figure 1A and B). Upon comparison with *Arabidopsis*, rice and maize homologs, the barley *HvNRPD/E2A* and *HvNRPD/E2B* genes exhibited 67% sequence identity to each other and they showed 56% identity to Arabidopsis *AtNRPD/E2.* The sequence similarity of *HvNRPD/E2A* is higher to the maize *NRPD2a* and *2b*, which are subunits of both POL IV and V (Haag et al., 2014), while the *NRPD/E2B* is more similar to the POL V-specific maize *NRPE2c* (Supplementary Figure 1C).

Utilizing the CRISPR/Cas9 system, two independent guide RNAs (sgRNAs) have been designed for each gene. In the T1 progenies we were able to identify a homozygous knock-out mutant for *NRPD/E2A* gene at the position of the second guide with 1 base pair (1-bp) deletion (*nrpd/e2a_1*, Figure 1A), while for the *NRPD/E2B* gene we detected 2-bp deletion at the position of the first guide RNA (*nrpd/e2b*, Figure 1A). To fully abrogate the RdDM pathway we also generated double knock-out mutants. In the T1 progeny we detected 1-bp insertion in *NRPD/E2A* gene at the sgRNA2 position and 1-bp deletion at the sgRNA3 position in *NRPD/E2B* gene creating the *nrpd/e2ab* double mutant (Figure 1A). These mutations are predicted to truncate the *NRPD/E2A* protein at domain 6 (*nrpd/e2a_1, nrpd/e2ab*), eliminating the majority of carboxyl terminus and *NRPD/E2B* protein at the domain 2 (*nrpd/e2b, nrpd/e2ab*) deleting the majority of protein coding sequences (Figure 1B). No mutations were detected at the other guide RNA positions. Additionally, we also identified two *nrpd/e2a* knock-out mutants bearing multiple mutations at the position of guide RNA2 (*nrpd/e2a_2*, *nrpd/e2a_3,* Supplementary Figure 1D).

**Figure 1.**
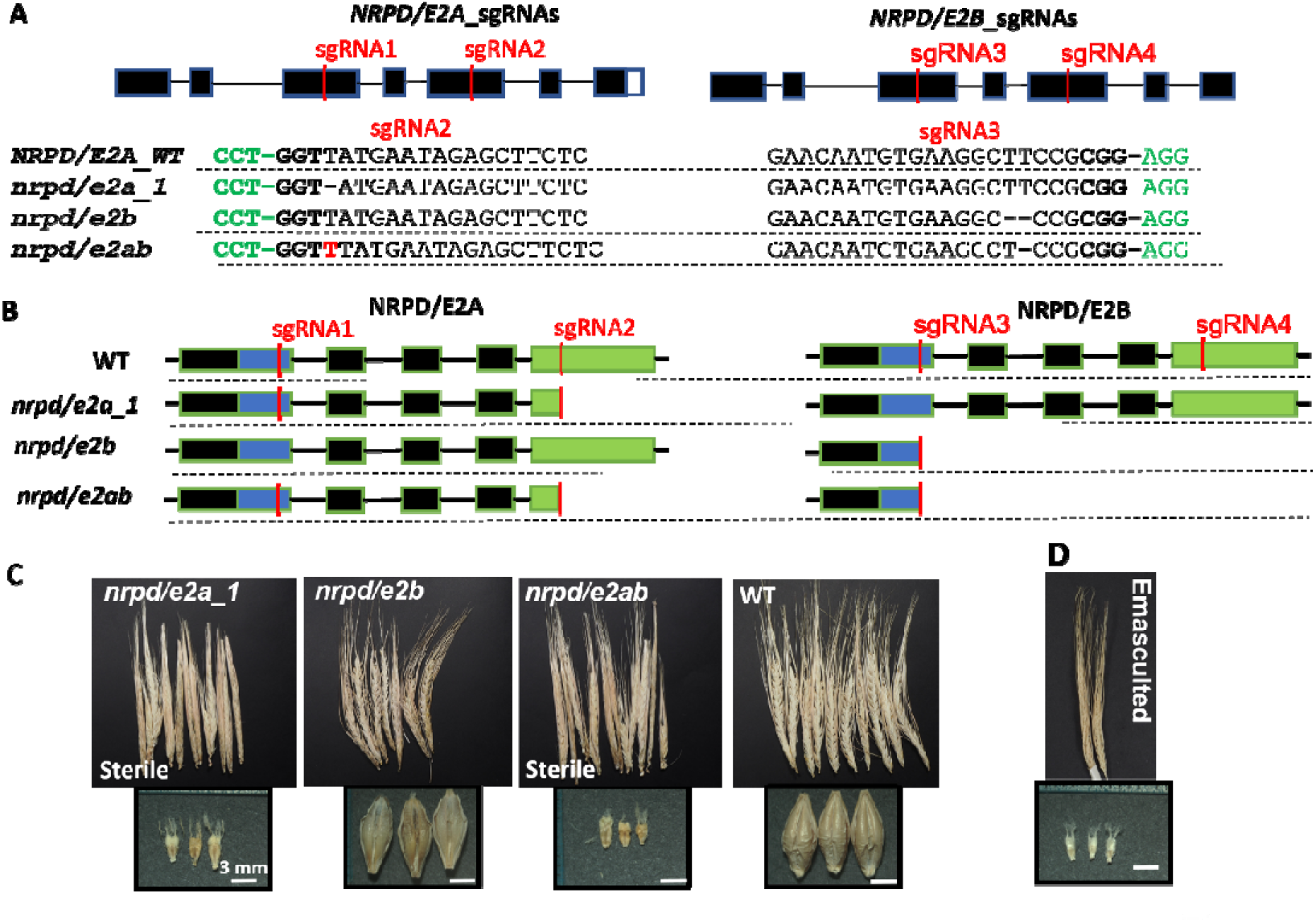
*nrpd/e2a* mutants exhibit full sterility. **A** Organization of barley *NRPD/E2A* (HORVU.MOREX.r3.2HG0206380) and *NRPD/E2B (*HORVU.MOREX.r3.7HG0706060) genes with positions sgRNAs (red lines) and the introduced mutations. Black boxes indicate exons while black lines show introns (upper panel). The dotted lines separate the genotypes of the mutant lines. Green letters indicate PAM sequences. *nrpd/e2a_1* and *nrpd/e2b* are single mutants having mutations in *NRPD/E2A* or *NRPD/E2B* gene, respectively*. nrpd/e2ab* is a double mutant bearing mutations in both, *NRPD/E2A* and *NRPD/E2B* genes (lower panel). **B** Representations of the protein domains (1-6) and the effects of introduced mutations. Blue box is the domain 2 and the site of sgRNA1/3. Green box is the hybrid binding domain (HBD) and the site of sgRNA2/4. Black boxes show all the other domains. The dotted lines separate the individual mutant lines. **C** Phenotypes of the spikes produced by T1 homozygous mutant lines and WT plants (upper panels). Dissected aborted caryopses of *nrpd/e2a_1* and *nrpd/e2ab* and mature seeds of *nrpd/e2b* and WT (bottom panels) **D** Phenotype of artificially emasculated spikes and dissected unfertilized ovary.

The mutant plants were grown in a greenhouse, under partially controlled environmental conditions, including temperature and pathogen management, and allowing for daily temperature fluctuations. Phenotypic analyses of the mutant lines revealed that the *nrpd/e2a* and *nrpd/e2ab* double mutants were completely sterile, yielding no harvestable seeds (Figure 1C and Supplementary Figure 1D). In contrast, the single *nrpd/e2b* mutant produced seeds with the size similar to that of the wild type plant (Figure 1C). This finding demonstrates that *NRPD/E2A* gene is a pivotal component of seed development and its loss-of-function induces complete sterility in barley.

A closer investigation of fully dried spikes from the *nrpd/e2a* and *nrpd/e2ab* mutants revealed the presence of aborted caryopsis ranging in size between 3-4 mm (Figure 1C, bottom panels). Notably, this size is significantly larger than that found in artificially emasculated spikes, which was measured to approximately 2 mm (Figure 1D). This difference strongly indicates that pollination and subsequent fertilization had successfully occurred and caryopsis development was arrested at approximately 3-5 days after pollination (DAP). Given that “Golden promise” is a strict self-pollinating cultivar, the initial development of these caryopses suggests that the mutant plants are capable of producing functional, viable pollen, pointing toward a post-fertilization developmental defect.

Altogether the analyses of several independent *nrpd/e2* mutants revealed that the presumed complete loss of RdDM activity in barley (driven by *nrpd/e2a* mutations) induces severe post-fertilization seed development anomalies, ultimately resulting in complete sterility under partially controlled greenhouse conditions.

### The barley *rdr2* mutants show partial abortion of seed development

Since the *nrpd/e2a* and *nrpd/e2ab* mutants were completely sterile, we were unable to propagate these lines for detailed downstream investigations in subsequent generations. To circumvent this and further dissect the impact of a disabled RdDM pathway on barley development, we generated mutant lines targeting the *RDR2* gene. RDR2 acts downstream of NRPD/E2 and utilizes POL IV-derived single-stranded transcripts as templates to synthesize complementary strands, thereby generating double-stranded RNAs (dsRNAs) that are subsequently processed by DCL3. Previously, we identified a single *RDR2* ortholog in barley (Hamar et al., 2020). Using CRISPR/Cas9 genome editing we produced three independent *rdr2* knock-out lines using two sgRNAs targeting distinct positions within the gene (Figure 2A). We identified single nucleotide insertions at the first and various deletions at second positions of the guide RNAs in three independent lines (*rdr2_1*, *rdr2_2* and *rdr2_3*) (Figure 2B). These frameshift mutations lead to premature stop codons, rendering the *RDR2* gene non-functional due to severe protein truncation (Figure 2C).

**Figure 2:**
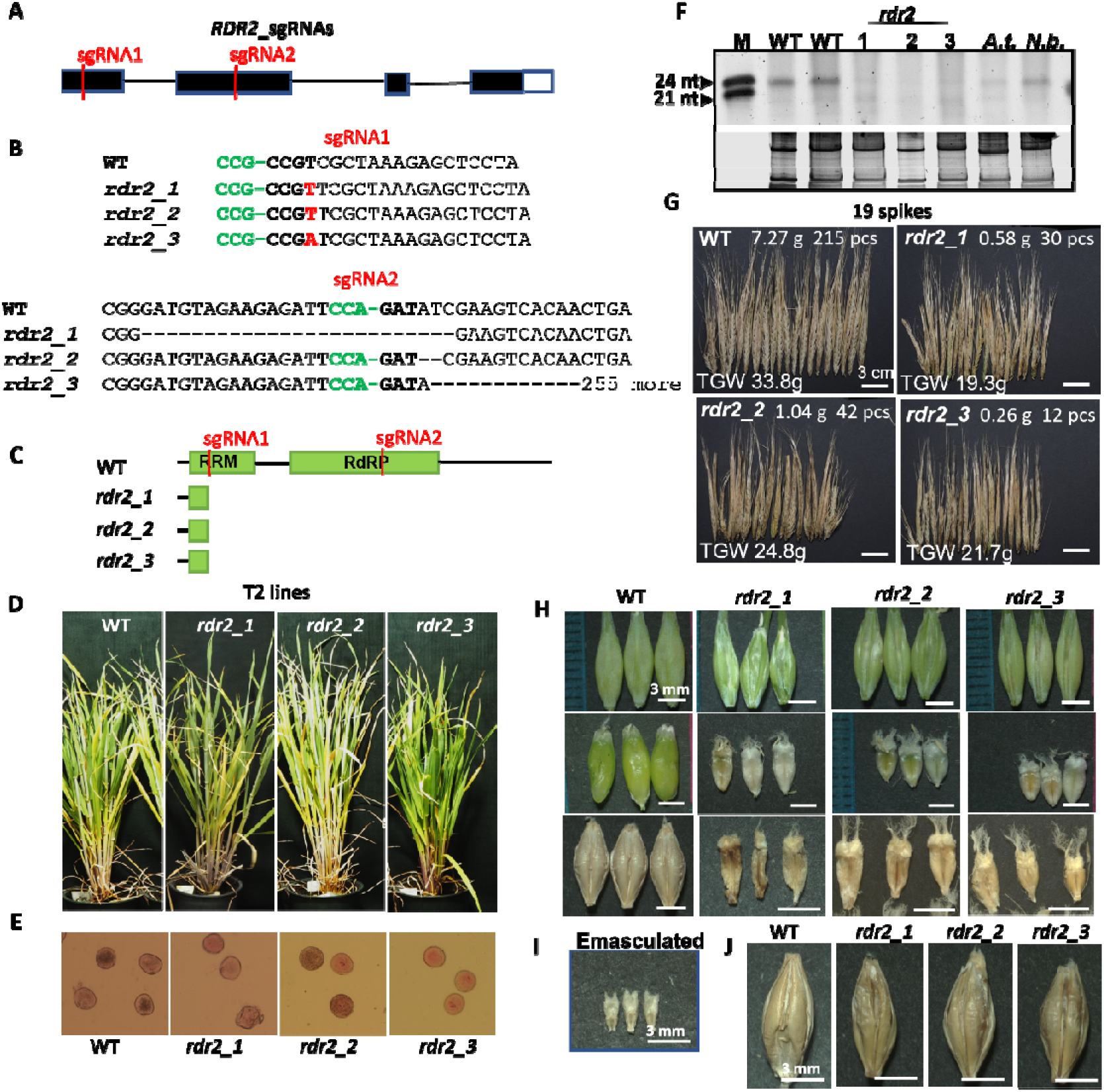
***rdr2* mutants exhibit partial sterility and seed development abnormalities A** Structure of barley *RDR2* (HORVU.MOREX.r3.2HG0174030) gene with the positions sgRNAs (red lines and letters), black boxes indicates exons and black lines introns, white box shows untranslated region. **B** Mutations generated mutations at sgRNA1 and sgRNA2 positions. **C** Schematic **r**epresentations of the effects of the introduced mutations. Green boxes show the domains. **D** Vegetative phenotype of mutant lines in T2 generation. **E** Stained pollens of mutant and WT pollens. **F** SYBR Gold staining of small RNAs of mutant, WT barley plants and control plants (*A.t.*; Arabidopsis, *N.b.*; *Nicotiana benthamiana*). Ribosomal RNAs serve as loading control (bottom panel). **G** Phenotypes of 19 spikes produced by homozygous mutant lines in T2 generation and WT plants. White numbers indicate weight, and number of the seeds and calculated thousand grain weight (TGW). **H** Green hulls of developing seeds (upper panels). Dissected developing green seeds (WT) and aborted green caryopses of mutant lines (middle panel). Dissected aborted dry caryopses (mutants) and mature seeds (WT) (bottom panels). **I** Phenotype of dried ovary in emasculated spike. **J** Phenotype of WT and mutant seeds.

The T1 generations of the mutant lines were grown in a partially controlled greenhouse environment with daily temperature fluctuations. The mutants exhibited similar vegetative development compared to the wild type (WT) plants (Supplementary Figure 2A). In contrast, the seed production was severely inhibited in the *rdr2* mutant lines. Mature spikes displayed a strongly reduced seed set with a production rate of approximately 12% relative to the WT, and the average mass of the harvested seeds was lower than that of WT seeds (Supplementary Figure 2B). Similarly, to the *nrpd/e2a* mutants, the development of the majority of caryopses was arrested at about 3-5 mm size range, at a post-pollination developmental step (Supplementary Figure 2C).

We further propagated the mutant lines to the T2 generation under identical greenhouse environment. The T2 plants of lines showed no drastic alterations in vegetative development (Figure 2D), although the mutants were typically slightly smaller in stature and showed delayed flowering compared to WT plants. According to pollen staining experiments, the mutants produced viable pollen with normal size and morphology (Figure 2E).

Since RDR2-derived dsRNAs serve as precursors for DCL3-mediated 24-nt siRNA biogenesis, *rdr2* mutants were expected to display a depleted accumulation of 24-nt siRNAs. We verified this by direct staining of total RNA extracts of young spikelets separated on polyacrylamide gels. In all three *rdr2* lines, the level of 24-nt siRNAs was strongly down-regulated compared to the WT plants, whereas signals representing the 21-nt small regulatory RNA population, (predominantly miRNAs) remained largely unaffected (Figure 2F).

The reproductive rates of the T2 generation closely mirrored those of the T1 plants, maintaining a drastic decrease in seed sets (reduced to 6-20% of WT levels) and a significant reduction in thousand grain weight (TGW) (Figure 2G). Examination of developing spikes confirmed the typical early arrest of caryopses at 3-5 DAP (Figure 2H), since they were significantly larger than the unpollinated ovaries of emasculated flowers (Figure 2I). Only a limited number of seeds managed to fully develop in mutant plants, and these mature seeds frequently exhibited morphological alterations (Figure 2J).

In summary, all the three independent *rdr2* mutant barley lines showed very similar phenotypic alterations such as reduced seed set due to early arrest of caryopsis development. The phenotype of *rdr2* plants was reminiscent of those of *nrpd/e2a* mutants but exhibited milder phenotype, since in contrast to *nrpd/e2a* lines, the *rdr2* mutants were able to produce mature seeds albeit with strongly reduced number and size. Taken together, our findings demonstrate that the RDR2-dependent branch of the RdDM pathway is a crucial component of seed development in barley.

### Loss of RDR2 activity triggers24-nt siRNA depletion and large-scale transcriptional reprogramming in caryopses

We performed high throughput small RNA-sequencing (sRNA-seq) and polyA-selected mRNA-sequencing (RNA-seq) analysis of developing mutant and wild type caryopses (Supplementary Figure 3A).

Analyses of sRNA reads revealed that the most abundant sRNAs in WT barley plants were the 24-nt long group (Figure 3A and Supplementary Figure 3F), which is a typical characteristic of plant sRNA profiles (Kasschau et al., 2007). Consistent with our northern blot data (Figure 2F), we observed a severe downregulation, amounting to a partial loss, of 24-nt long siRNAs across all three *rdr2* mutant lines. Additionally, subtle alterations were detectable in the abundance of 21-, 22-and 23-nt long size classes in the *rdr2* mutants. Next, we defined the 24-nt siRNA producing genomic clusters on the barley genome, identifying 2357 clusters, which have higher expression than 1 RPM (reads per million) in at least one sample group (WT or the three mutants) (Supplementary Table 1). The principal component analysis (PCA) of the clusters clearly segregated the mutant lines from the WT plants (Supplementary Figure 3D). Notably, we identified 979 commonly downregulated 24-nt siRNA clusters in all *rdr2* mutants, representing 60.8 % of the expressed clusters in WT, with at least 10-fold reduction compared to WT caryopses (Figure 3B and Supplementary Table 1). This strong depletion indicates that these 24-nt siRNAs are direct products of RDR2. The number of downregulated clusters exclusive to each mutant line were low, reinforcing the highly consistent molecular phenotype among the independent mutants. While a limited number of upregulated 24-nt producing clusters were identified in the *rdr2* mutants, only 27 were shared among all three lines (1.7% of the WT clusters) (Figure 3C). Together, these data demonstrate that mutations in the barley *RDR2* gene globally impair the activities of 24-nt siRNA generating genomic clusters.

**Figure 3.**
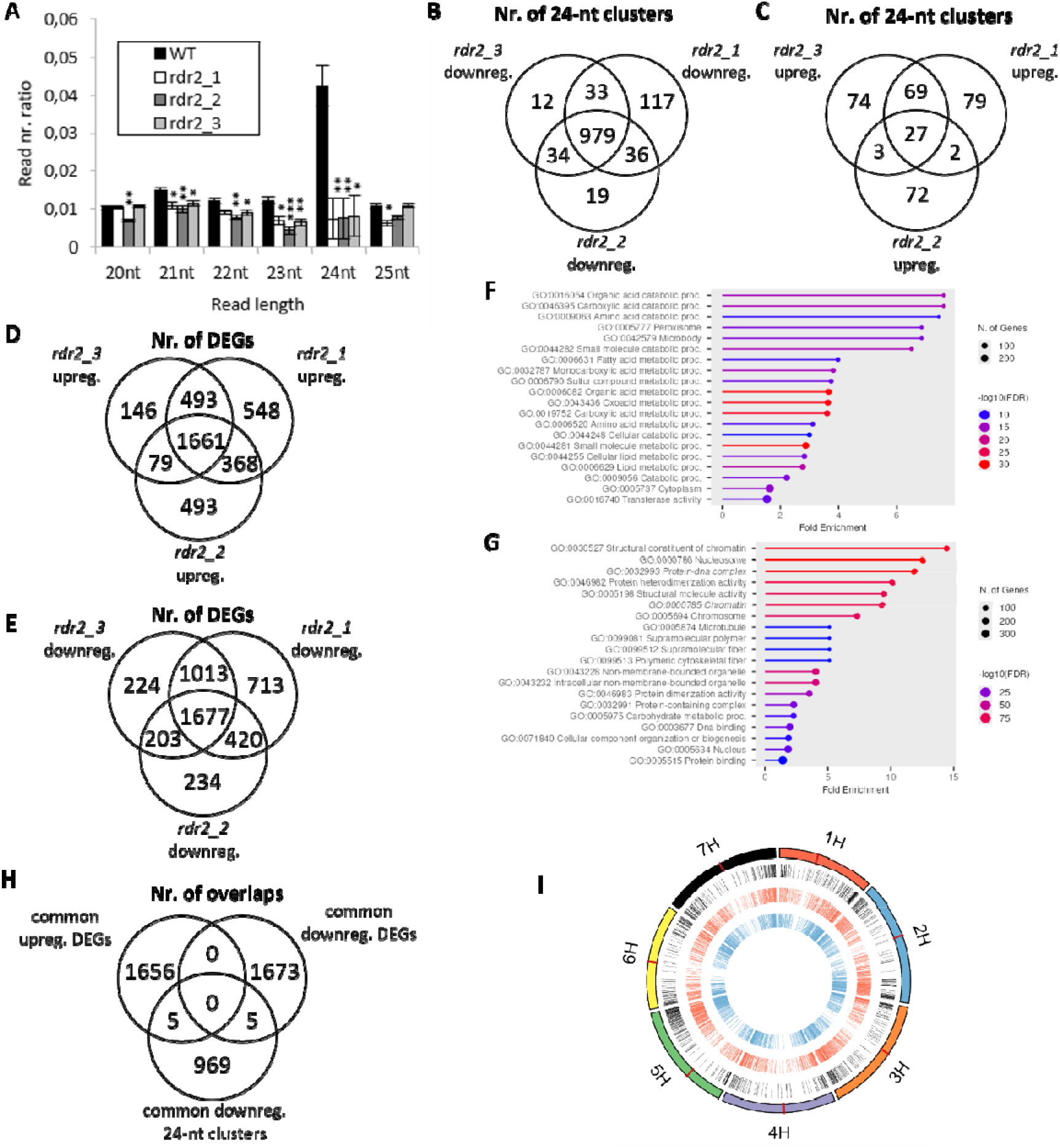
sRNA-and poly(A)-selected RNA-sequencing of *rdr2* mutant caryopses. **A** Ratio of reads with 20-25 nt in the small RNA sequencing data of wild type (WT) and *rdr2-1*, *rdr2-2*, *rdr2-3* mutant caryopses. Bars represent standard errors, n ≥ 2, *P*-values are calculated for mutants vs WT data based on 2-tailed Student’s *t*-test (\**P* < 0.05, \*\**P* < 0.01, \*\*\**P* < 0.001). **B-C** Venn diagrams with the number of down-(B) and upregulated (C) 24-nt RNA clusters in the mutants compared to WT caryopses. **D-E** Venn diagrams with the number of up-(D) and downregulated (E) differentially expressed genes (DEGs) in the mutants compared to WT. **F-G** 20 most significant gene ontology terms of up-(F) and downregulated (G) DEGs. **H** Venn diagram with the number of commonly downregulated 24-nt RNA clusters and up-or downregulated DEGs including their promoter region (up to-2000 bp to the transcriptional start) in the mutants. Intersections represent the number of overlapping genomic regions. **I** Schematic representation of the seven barley chromosomes (outer circle) and the commonly downregulated 24-nt RNA clusters (black), up-(red) and downregulated (blue) DEGs aligned to them. Centromeres are represented with dark red lines.

Parallel mRNA-seq analyses were conducted to evaluate the transcriptome of caryopses samples. The PCA showed a distinct separation between the mutant and WT sample groups (Supplementary Figure 3B). We observed profound changes in the expression of genes in the mutants relative to the WT plants, identifying 1661 common significantly up and 1677 downregulated genes in *rdr2* mutants with at least two-fold change (Figure 3D and E, Supplementary Table 1). A smaller but still notable fraction of differentially expressed genes (DEGs) was specific to individual mutant lines, indicating that besides the core transcriptomic response, individual lines can show minor variations. These differences can be attributed to potential sampling variations or line-specific alteration in epigenetic changes. Gene Ontology (GO) term enrichment analysis revealed that the commonly upregulated genes predominantly associated with catabolic processes (Figure 3F), while the downregulated genes mainly belonged to chromatin associated functions, many of which are histone proteins (Figure 3G). Finally, we cross-referenced the genomic coordinates of the 979 downregulated 24-nt RNA producing clusters with the positions of up-or downregulated genes and their promoters (defined as 2000 nt upstream of the transcription start site (TSS)). Only marginal spatial overlaps were detected between these regions (Figure 3H).

To better understand their spatial relationship, we mapped the genomic localization of the downregulated 24-nt siRNA clusters and the up-and downregulated genes on the seven barley chromosomes. In contrast to findings in *Arabidopsis* (Karányi et al., 2022), the density of siRNA producing clusters was significantly higher towards the distal (telomeric) regions of the chromosomes (Figure 3I). Both the up-and downregulated genes exhibited a highly similar distribution pattern along the chromosome arms. Since 24-nt siRNAs typically act in *cis,* this observation suggests a spatial regulatory correlation between siRNA producing loci and the affected genes. The finding of very limited number of 24-nt siRNA direct hits in gene bodies or promoters of the affected genes suggests that these 24-nt siRNAs affect the gene expression landscape through indirect epigenetic/heterochromatic reorganization or relaxation, rather than local *cis*-primed RdDM silencing of specific protein-coding genes.

In conclusion, our transcriptome data confirm the partial loss of 24-nt siRNAs in all the *rdr2* mutants and reveals a widespread transcriptome reprogramming in barley caryopsis. The RdDM mediated regulation may predominantly act indirectly on gene expression regulation rather than direct, localized gene targeting.

### *rdr2* mutant exhibits endosperm developmental defects and inhibited germination

Given the highly consistent phenotypes among all three independent lines, the *rdr2-2* mutant was selected for in-depth characterization using T3 generations. Under strictly controlled conditions ensuring the proper temperature level (not higher than 21 °C) and pathogen free environment. WT plants exhibited enhanced seed production compared to previous trials (Figure 2G), while the performance of the mutant *rdr2_2* drastically increased. We found that the seed production capability of *rdr2_2* reached cca 50% of the wild type plants in numbers, in contrast to less controlled conditions (Figure 4A and Supplementary Figure 4A). This environmental responsiveness suggests that the mutant line possesses heightened stress sensitivity. Despite the partial rescue, the developing spike of the *rdr2*_*2* line exhibited various morphological distortions (Figure 4A and Supplementary Figure 4A), the size of the seeds were smaller and of abnormal shape (Figure 4B) and their germination rates were reduced to about 30% relative to the WT seeds (Figure 4C and Supplementary Figure 4B).

**Figure 4:**
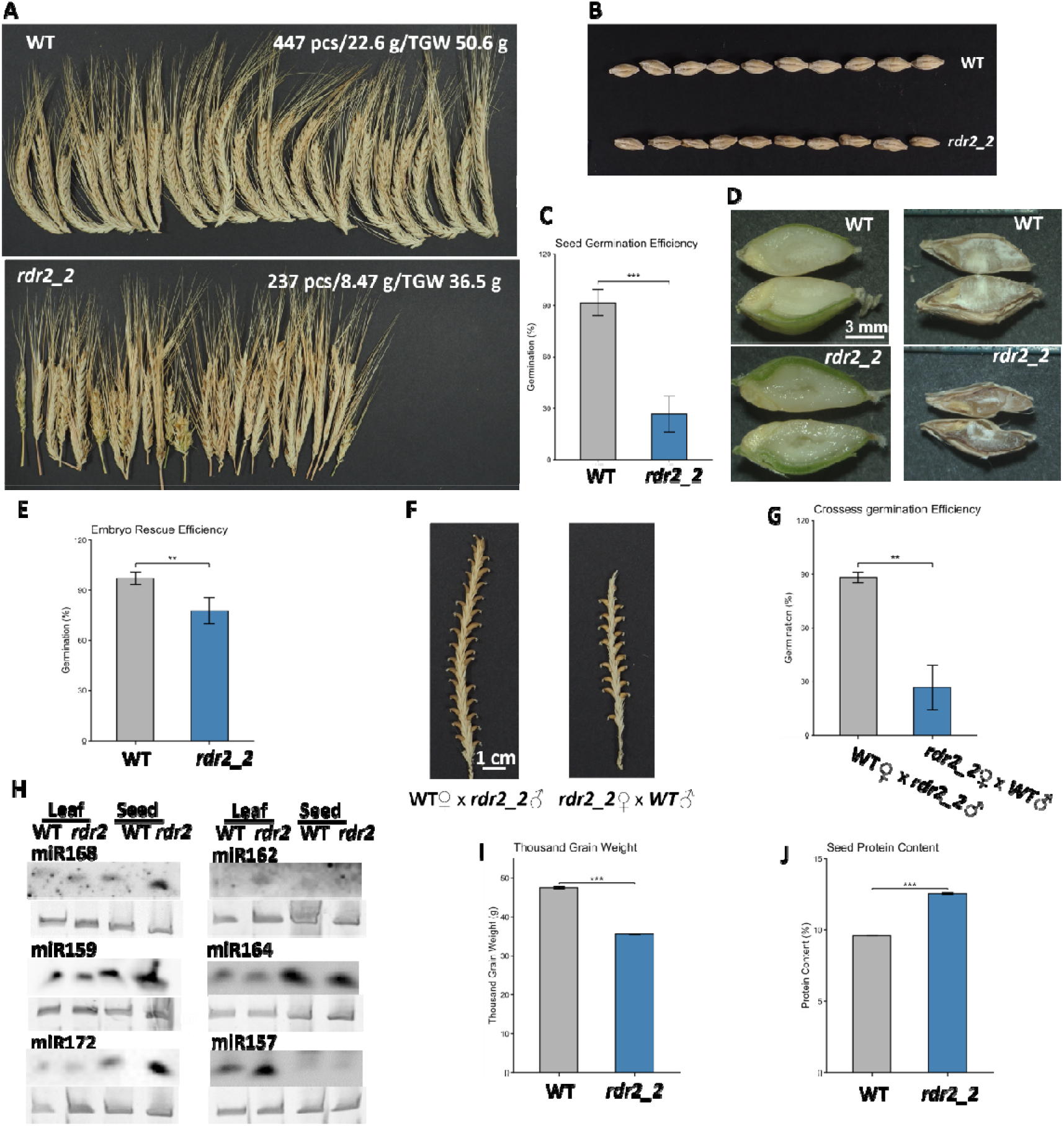
***rdr2_2* shows endosperm abnormalities and seed defects under controlled condition. A** 25 representative spikes of wild type (WT) and mutant (*rdr2_2*) plants, white text on the top right shows number of seeds from the 25 spikes, weight of the seeds and thousand grain weight (TGW) respectively. **B** seed morphology of the WT and *rdr2_2* plants. **C** Seed germination assay (%) of WT and *rdr2_2*. **D** Dissected green seed (Left) and dried seed (right) of WT and *rdr2_2*. **E** Efficiencies of embryo rescue experiments (%) of WT and *rdr2_2*. **F** Crossing experiments, WT mother crossed with *rdr2_2* pollen (left) and *rdr2_2* mother crossed with WT pollen. **G** Germination efficiencies (%) of seeds deriving from crossing experiments. **H** Micro RNA northern blots of WT and *rdr2_2* from leaf and seed tissues. Bottom panels show EtBr stained rRNAs as loading controls. **I-J** Thousand grain weight (g) (**I**) and seed protein content (%) (**J**) of WT and *rdr2_2*, respectively. Data are mean ± SD of three independent replicates (*, p<0.05; **, p<0.01; ***, p<0.001 by student’s t test).

To determine the cause of the poor germination, we investigated the endosperm formation of barley WT and *rdr2_2* mutant seeds. Longitudinal sections revealed that in contrast to the homogenous structure of the WT endosperm, the mutant endosperms have prominent internal cavities (Figure 4D and Supplementary Figure 4C). In line with this observation the dried mature mutant seeds show reduced mass of endosperm content, about 28% reduction in TGW (Figure 4A and 4I). The cross sections also revealed that the embryos show normal external phenotype (Figure 4D and Supplementary Figure 4C). To test whether the poor germination ability of the mutant is due to defects of the embryo or the endosperm, we carried out embryo rescue experiments. Green immature seeds were harvested from *rdr2*_*2* and WT plants and embryos were dissected and moved to artificial growing medium. WT immature embryos exhibited close to a hundred percent (95 %) rate of successful regeneration to healthy seedlings and later fully developed normal plants. The *rdr2*_*2* embryos also showed high albeit slightly less regeneration efficiency reaching about 80% (Figure 4E and Supplementary Figure 5). This finding indicates that the embryo development of the *rdr2_2* is only moderately affected, strongly suggesting that germination failure is primarily driven by the defective endosperm.

In *Brassica rapa,* RdDM mutants are also associated with seed abortions and this trait was controlled by the maternal sporophytic genotype(Grover et al., 2018). To test this inheritance pattern in barley, we performed reciprocal crosses between WT and *rdr2_2* plants. Artificial pollination of WT mother plants with *rdr2_2* pollens was highly efficient with nearly all flowers setting seeds (Supplementary Figure 4D). Pollination of *rdr2_2* mother plant with WT pollen was able to induce seed development with less efficiency, approximately 70% percent of the pollinated flowers produced seeds (Figure 4F, Supplementary Figure 4D) suggesting that seed abortion is mainly controlled by the maternal sporophytic genotype. Next, we tested the germination efficiency of the seeds produced during the crossing experiments. We found the seeds derived from WT♀ x *rdr2_2*♂ germinated very efficiently (∼90%), while *rdr2_2*♀ x WT♂ seeds germinated poorly, similarly to the self-pollinated *rdr2_2* plants (∼30%) (Figure 4G, Supplementary Figure 4E). These genetic data confirm that partial seed abortion and germination inhibition are predominantly determined by the maternal sporophytic genotype.

Next we monitored the accumulation of selected conserved miRNAs. Small RNA northern blot analyses of total RNA samples extracted from *rdr2_2* leaves and immature green seeds showed comparable or in some specific cases higher signal than WT for the investigated miRNAs, indicating that disruption of RdDM may impact miRNA biogenesis indirectly only in certain cases (Figure 4H).

Detailed metabolic/biochemical investigation of grain parameters also revealed drastic change in TGW value which was accompanied by significant increase in relative protein content in the mutant seeds (Figure 4I and 4J). This strong negative correlation suggests that the higher protein percent in the *rdr2_2* seeds is a concentration effect resulting from the decreased starch accumulation and endosperm mass. The starch composition analyses show very similar approx. 28% amylose + 72% amylopectin ratio in the mutant and WT plants (Supplementary Table 2). However, the protein composition shifted significantly due to the mutation, characterized by an increased ratio of monomeric prolamins over glutelins. Furthermore, since protein composition varied with grain size, which supports a compensatory mechanism, the reduction of specific structural proteins directly impacts final grain mass while triggering the synthesis of alternative protein fractions. Moreover, the smaller mutant seeds also have a larger relative surface area consequently the proportion of cell wall-forming components in the husk increases indicated by the higher β-glucan content (Supplementary Table 2). Altogether these data support that *rdr2_2* mutant exhibits inhibited germination efficiency likely due to endosperm development anomalies.

To reveal changes at molecular level, we dissected endosperm samples from green seeds and carried out sRNA-seq and polyA-selected RNA-seq analysis. sRNA-seq analyses, in line with the caryopsis data, revealed drastic reduction of 24-nt siRNA amount in the endosperm of *rdr2_2* mutant (Figure 5A, Supplementary Figure 3G). Intriguingly, we also observed a severe downregulation in the reads of other size sRNAs, including 21-nt long sRNAs, typically miRNAs. This finding indicates that in endosperm the production of sRNAs is affected in general by the inhibition of RdDM pathway suggesting a crosstalk between the RNAi pathways.

**Figure 5:**
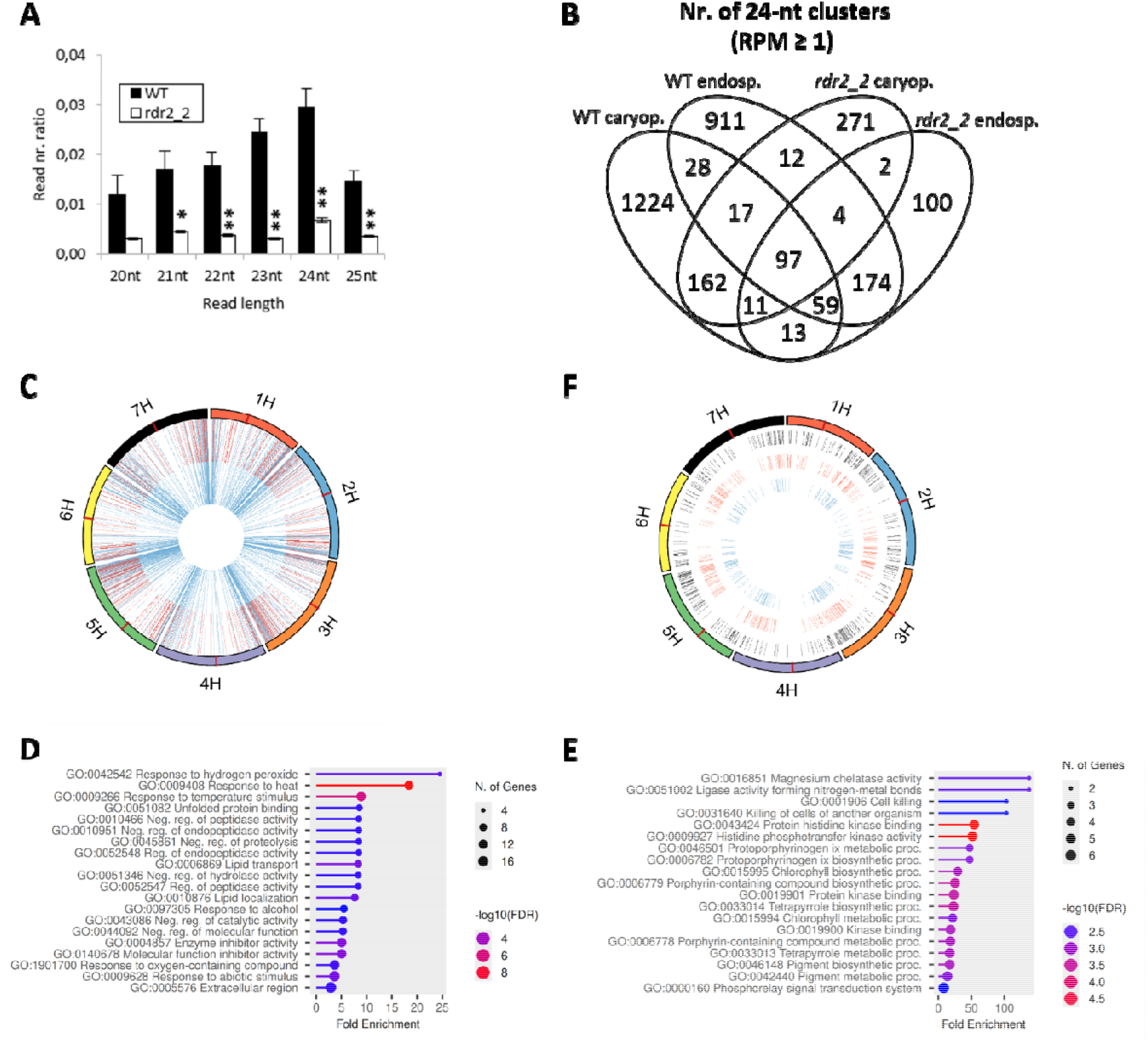
sRNA-and poly(A)-selected RNA-sequencing of *rdr2_2* mutant endosperm and heat stress sensitivity of *rdr2_2*. **A** Ratio of reads with 20-25 nt in the small RNA sequencing data of wild type (WT) and *rdr2-2* mutant endosperm. Bars represent standard errors, n = 3, *P*-values are calculated for mutant vs WT data based on 2-tailed Student’s *t*-test ( \**P* < 0.05, **\*\****P* < 0.01, **\*\*\****P* < 0.001). **B** Venn diagram with the number 24-nt RNA clusters with at least 1 RPM (read per million) in the *rdr2_2* and WT caryopses and endosperm. **C** Schematic representation of the seven barley chromosomes (outer circle) and the downregulated 24-nt RNA clusters aligned to them in caryopses (red) and endosperm (blue). Centromeres are represented with dark red lines. **D-E** 20 most significant gene ontology terms of up-(D) and downregulated (E) DEGs in *rdr2_2* endosperm. **F** Schematic representation of the seven barley chromosomes (outer circle) and the commonly downregulated 24-nt RNA clusters (black), up-(red) and downregulated (blue) DEGs in endosperm aligned to them. Centromeres are represented with dark red lines

Detailed analysis of the comparison of all *rdr2_2* and WT caryopsis and endosperm 24-nt producing clusters, showed extensive spatial dynamics. In WT tissues, we identified a highly tissue-specific landscape, with 1,410 clusters exclusive to the caryopsis, 1,101 exclusive to the endosperm, and only 201 shared between them (Figure 5B, Supplementary Table 1). In the *rdr2_2* caryopsis and endosperm the number of the clusters strongly dropped (to 576 in caryopsis and 574 in endosperm). Moreover, the clusters active in *rdr2_2* mutants showed only partial overlaps with their WT counterparts, since 289 clusters in caryopsis and 126 clusters in endosperm did not show overlaps with the WT samples (Figure 5B), indicating that 24-nt siRNA producing clusters were activated only in the mutants. Chromosomal mapping verified this tissue-specific divergence (Figure 5C), and confirmed that the remaining downregulated 24-nt siRNA clusters maintain a preferential accumulation near the telomeric regions on all seven barley chromosomes (Figure 5C). Finally, RNA-seq analyses of the *rdr2-2* endosperm revealed gene expression changes of lower magnitude than those found in the caryopsis, identifying 345 significantly up-and 175 downregulated genes (Supplementary Table 1). GO term enrichment analysis revealed that the upregulated genes are mainly associated with stress responses, particularly heat stress responses (Figure 5D), while downregulated genes typically connected to metabolic processes, consistent with the anomalies of endosperm development (Figure 5E). Localization of the downregulated 24-nt siRNA producing clusters and the up-and downregulated genes on the seven chromosomes, similarly to caryopsis, showed a distribution with higher density closer to the termini of chromosomes (Figure 5F). These findings further support the role of RdDM pathway in gene expression regulation via chromatin structure alterations.

### Long term heat stress memory in barley is controlled by RdDM

Previously we have demonstrated the heat-stress inducible nature of barley RdDM pathway, by showing key components such as HvDCL3, HvRDR2, HvRDR6 and HvAGO6 exhibit increased expression under heat stress (HS) conditions (Hamar et al., 2020). Moreover, our recent transcriptomic data revealed that *rdr2-2* seed production fluctuates depending on growing conditions, especially temperature variation, and HS-related genes are activated in the endosperm. These observations prompted us to directly investigate the roles of RDR2 in HS-sensitivity and thermo memory. For this we followed the previously established thermotolerance memory protocol (Pratx et al., 2025), and evaluated the phenotypic outcomes, focusing on plant survival and growth rate (Figure 6A and Supplementary Figure 6). Under ambient growth conditions the non-treated *rdr2_2* plants were inherently shorter than the WT plants (Figure 6B). The basal heat stress (HS only) treatment typically killed most of the plants. Although primed (acclimated, ACC) *rdr2-2* mutants successfully survived the lethal HS challenge similarly to WT plants, they exhibited a significantly more pronounced relative growth reduction compared to non-treated controls and WT plants (Figure 6B and Supplementary Figure 6). This finding indicates that the inhibition of RdDM pathway interferes with the long-term HS memory, rendering the ACC plant more vulnerable to HS.

**Figure 6:**
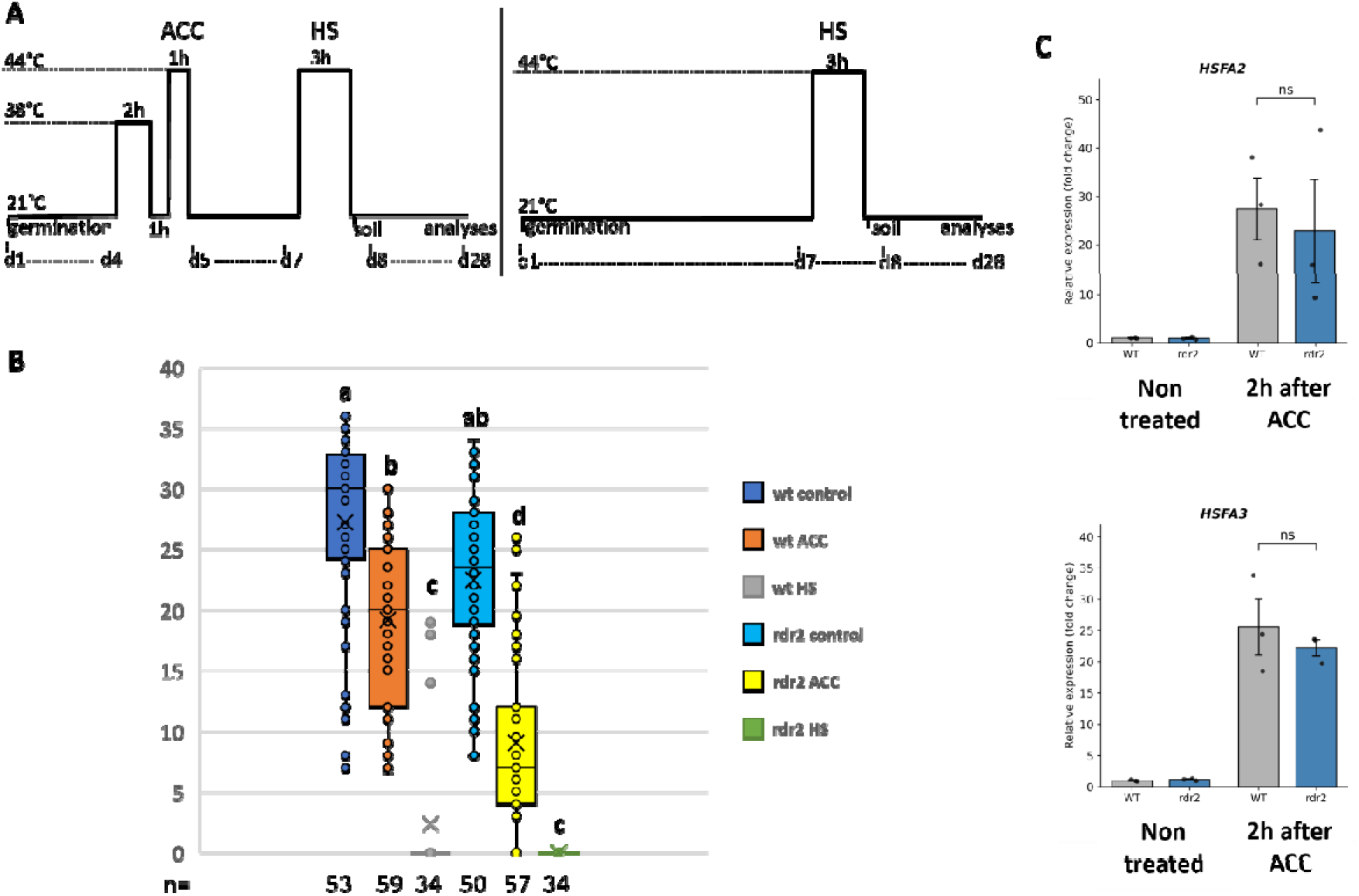
A *rdr2* mutant exhibits reduced heat stress memory capacity. **A** Heat stress (HS) treatment scheme. Plants were two-step acclimated (ACC) and exposed to heat stress as indicated (left panel). HS control plants were exposed to heat stress without acclimation (right panel). Control plants were not treated. h=hour, d=day. **B** Heights of WT and *rdr2_2* plants after 20 days of HS treatment were measured and compared to control plants. ACC, HS only and non-treated control plants. Data of three independent biological replica experiments were summarized in the box plot. Significantly different datasets were labelled with different letters based on one-way ANOVA test. **C** Transcript levels of *HSFA2* (upper panel) and *HSFA3 (*bottom panel) in WT (gray) and *rdr2_2* (blue) under control condition and 2h after ACC. WT, *rdr2_2*. Data are mean ± SEM of three replicates. ns= not significant according to Welch’s t-test comparison of WT and *rdr2_2*.

To test if RdDM mediated HS tolerance is achieved via HSFA2 and/or HSFA3 pathways (Pratx et al., 2025), we monitored the expression profile of these genes. RT-qPCR revealed no significant alterations in their expression in *rdr2_2* compared to the WT plants, under either non-treated or two hours after ACC treatment (Figure 6C). The unaltered expression of *HSFA2* and *HSFA3* genes indicates that RDR2-dependent thermotolerance operates independently of canonical HSF signaling, likely functioning through a chromatin-mediated epigenetic mechanism.

## Discussion

In the present work we identified barley *NRPD/E2* genes and created *NRPD/E2* and *RDR2* CRISPR mutants to show these are indispensable for proper caryopsis/ seed development. To unravel the underlying molecular and cellular changes, we integrated RNAseq and sRNAseq transcriptome profiling with histological/microscopic observation and seed content measurements.

Barley, common wheat (*Triticum aestivum*) and rye (*Secale cereale*) are closely related and are the members of the *Triticeae* tribe. The barley genome size is large, 5.1 Gbp with >80% of repetitive elements (Mayer et al., 2012). The individual sub-genomes of hexaploid wheat exhibit the same genetic composition as barley in terms of the genome size, gene content, and repetitive element contents (Middleton et al., 2013). Consequently, despite its large genome size, the diploid nature of barley makes it the best genomic model for hexaploid wheat. Given the increased TE content of the *Triticaea* genome, the regulatory importance of the RdDM pathway likely expands beyond its canonical roles described in small-genome models, taking on broader epigenetic governance over transcription.

The importance of the RdDM pathway in barley was highlighted by the phenotype of mutant plants bearing mutations in *NRPD/E2* and *RDR2* genes. These homozygous mutants were able to grow and develop flowers, spikes and fertile pollen, but show an early abortion during caryopsis development. Under our growing conditions, the *nrpd/e2a* mutants showed full sterility and no seeds were produced. This finding is in line with rice data, where null alleles of NRPD/E2 also show sterility (Chakraborty et al., 2022). The barley *rdr2* mutants also showed an early developmental arrest of caryopses, but the spikes, showing abnormal growth and occasional branching, were able to produce seeds although morphologically aberrant, having inhibited germination properties. The milder phenotype associated with *rdr2* mutants can be the outcome of redundant background activities of other RDR proteins. The reduced fertility phenotype was described for other various RdDM rice mutant plants (Moritoh et al., 2012; Wei et al., 2014; Xu et al., 2020; Zheng et al., 2021). We also experienced that the seeds produced by *rdr2* mutants were smaller in size and showed deformation compared to the wild type seeds probably due to reduction of the endosperm mass and formation of caveats in the structure of the endosperm. The underlying changes in histone and chromatin structure concurrently restricted cell division, resulting in smaller and lighter grains, while reprogramming the expression patterns of seed storage proteins.

Since the peak synthesis of monomeric prolamins (gliadins) and glutelins occurs at different stages during grain development, variations in grain size and weight directly determine the time and energy allocated for the accumulation of each protein group. If the synthesis of certain proteins, such as the macromolecular matrix-forming glutelins, is reduced due to mutation or environmental impacts, the plant attempts to counterbalance this nitrogen and energy deficit by enhancing the production of other proteins, such as monomeric prolamins. The extent of this compensation is directly reflected in the final grain weight. Furthermore, this significant correlation indicates that the mutation did not merely affect protein genes, but globally altered the developmental program of the endosperm (Xhaferaj et al., 2023; Marín-Sanz et al., 2026).

Similarly, nutrient filling defects of endosperm were described in rice mutated in chromatin remodeler OsCLSY4, a major upstream regulator of the RdDM pathway (Pal et al., 2025; Zhang et al., 2026). The *rdr2* mutants produced seeds showing inhibited germination, which can be connected to endosperm development problems. Altogether, these observations suggest that regardless of the genome size (rice has a relatively small genome of 430 Mb), the control of RdDM on reproductive development can be deeply conserved in modern starch grains.

In rice the disruption or inhibition of the RdDM pathway is often associated with severe vegetative and reproductive abnormalities such as semi-dwarfed stature, reductions in tiller number, delayed heading or no heading, abnormal panicle and spikelet morphology (Moritoh et al., 2012; Wei et al., 2014; Xu et al., 2020; Zheng et al., 2021; Pal et al., 2025; Zhang et al., 2026). In contrast, we found that stature of *rdr2* plants was quite similar to that of WT plants, although the fully developed mutant plants are slightly smaller, no severe dwarfism was detected. The average spike number per plant of the mutant *rdr2* is also similar to the WT and the mutant plants show about a two-week delay of flowering. These findings indicate that rice and barley share common RdDM functions but also show differences.

RdDM predominantly targets TEs and TE fragments positioned adjacent to the genes within transcriptionally permissive euchromatic regions (Deniz et al., 2019). During early caryopsis development (3-5 DAP) failure of RdDM may trigger local epigenetic instability resulting in TE reactivation and perhaps chromosomal instability (Wen et al., 2026), rather than a direct RdDM-derived silencing of specific protein-coding genes. We observed enrichment of catabolic-and stress-related transcripts, alongside a severe downregulation of histone transcripts in *rdr2* mutants. Histone gene transcription is almost certainly a consequence rather than cause: histone production is strictly coupled with DNA replication in the cell cycle S phase. In case of genomic stress and cell division arrest, histone transcription is promptly arrested to prevent accumulation of a massive amount of positively charged proteins within the cell (Raynaud et al., 2014). During the early period of embryo development (3-5 DAP) the fault in RdDM activity may cause a heterochromatic decondensation that results in transposon bursting and chromosome fragility. The physical cavities of endosperm we observe in mutant seeds, could be caused by (i) localized nuclear death, a subset of nuclei with the syncytium suffers severe chromosomal damage due to transposon activation/mis-regulation of chromatin structure which can be associated with apoptosis, leaving physical gaps, and (ii) failure of transition from syncytium to cellular tissue within the endosperm (catabolic processes are down regulated, therefore normal cell wall deposition absence could leave holes behind). The embryo which is entirely dependent on the endosperm will starve and finally arrest its development. This idea is supported by our embryo rescue assays. Imprinting crisis (parent of origin) may be also an important regulatory factor. Imbalance in maternal vs paternal gene dosage universally triggers early seed abortion and embryo arrest. This idea is backed by our backcross experiment. In summary, we propose the following possible cascade of events: (a) loss of RdDM enzymes NRPD/E2 and RDR2 functions lead to depletion of het-siRNAs; (b) heterochromatic DNA and TE are de-repressed; (c) this causes mitotic and chromatin organization stress; (d) leading to global endosperm cell cycle arrest and alteration of overall transcriptome, including histone gene repression; (e) next syncytial nuclei either die or cavities are formed within the developing endosperm tissue, and (f) ultimately caryopsis abortion or embryo starvation occur.

In conclusion our findings show that RdDM appears essential for vegetative HS acclimation in barley, as its disruption in *rdr2_2* mutant results in inhibited acclimation process rendering the mutants more vulnerable to subsequent heat stresses. Our data indicate that RdDM is a pivotal regulator of both reproductive development and stress responses in barley. The similarity of RdDM functions in seed set control and endosperm development and the differences in the growth regulation between rice and barley highlight a possible common origin of cereal RdDM function, that underwent lineage-specific sub-functionalization for stress adaptation. Intriguingly, in dicots different phenotypes associated with the lack of RdDM activity (Wang et al., 2020; Dew-Budd et al., 2024; Feng et al., 2025) indicating the evolutional flexibility of the system. From a biotechnological perspective, this feature may be useful, allowing the targeted modulation of RdDM pathways to achieve enhanced agronomic performance. Therefore, the thorough and detailed exploration of RdDM roles in nurturing embryo development processes and heat stress responses need to be tasks for future research work holding numerous points of interest and value for technological perspectives, as well.

## Materials and Methods

### Plant materials and growth conditions

Barley (*Hordeum vulgare*) ‘Golden Promise’ wild type and genome edited seeds were sown into Jiffy-7 (44 mm) Substrate Pellet and the developing young plantlets were grown in growth cabinet (Versatile Environmental Test Chamber MLR-350; Sanyo, Tokyo, Japan) under 18 °C daytime and 12 °C night temperatures with 16-h light and 8-h dark periods. After 3-4 weeks the plants were transferred into pots and further grown either in semi-controlled greenhouse conditions, allowing significant daily temperature and light intensity fluctuations or in plant growing room under constant 20°C with a photoperiod of 16-h light (400 µmol/m^2^/s) and 8-h dark.

### Generation of barley RdDM CRISPR/Cas9 mutants

We used the genomic sequences of the target genes for designing sgRNAs suitable for generating knock-out genome edited lines for a *NRPD/E2A* (HORVU.GOLDEN_PROMISE.PROJ.2HG00186300 and HORVU.MOREX.r3.2HG0206380), *NRPD/E2B* (HORVU.GOLDEN_PROMISE.PROJ.7HG00631350 and HORVU.MOREX.r3.7HG0706060) and *RDR2*( HORVU.GOLDEN_PROMISE.PROJ.2HG00155150, HORVU.MOREX.r3.2HG0174030). CRISPOR software (Concordet and Haeussler, 2018) was used to obtain CRISPR/Cas9 target sites on important (domain 2 and domain 6 of the NRPD/E2) protein domains and sgRNAs with minimal off-target activities were obtained. For each of the two *NRPD/E2* genes, two sgRNAs were selected and construct was made based on the tRNA-processing system (Xie and Yang, 2019) and the four sgRNAs were combined for generation of double mutants of the *NRPD/E2A* and *NRPD/E2B* using the same system. For the *RDR2*, two sgRNAs were obtained also targeting important (RNA recognition motif (RRM) and RNA dependent RNA polymerase (RdRP)) domains (Supplementary Table 3). The constructs were created using the multiplex approach described previously (Xing et al., 2014). For each*, Agrobacterium* mediated method was used to transform immature embryos and selection was carried out using hygromycin supplemented medium (Kis et al., 2019). Mutants were first identified by screening using T7 endonuclease assay and the sequence composition of the mutations were determined by Sanger sequencing.

### Germination assay, embryo rescue and artificial pollination

To assess the germination efficiency of different genotypes, wild type and mutants, seeds were placed onto two layers of wet filter papers layered on wet cotton wool in closed plastic boxes. Sterile distilled water was used to wet the filter papers and seeds were germinated at 20-22°C for 3 days in dark and 4 days in the light. Each experiment was repeated three times independently.

To rescue the embryo, green matured seeds were collected from mother plants, surface sterilized and the embryo extracted under microscope in a laminar hood chamber. The extracted embryos were grown on medium (MS supplemented with maltose, glutamine, myoinositol, ammonium nitrate and thiamine-HCL with Gelrite as a gelling agent) at 18-19°C. Each experiment was repeated three times.

Pollinations were carried out on potted barley plants grown under greenhouse conditions. Anthers were removed from the mother florets 2–3 days before the anthesis using tweezers. Pollination was performed 3 days after emasculation using a barley spike in the anthesis stage.

### Northern blotting and small RNA staining

Frozen samples were first grinded in mortar with liquid nitrogen, extraction buffer (0.1 M glycine-NaOH, PH 9.0, 100 mM NaCl, 10 mM EDTA, 2% SDS) was added and the standard Phenol/Chloroform method was followed. For the microRNA analysis, 6µg of total RNA was run on 12% polyacrylamide gels with 8 M urea, transferred to Hybond NX membrane (GE Healthcare, Chicago, Illinois USA) with semi-dry blotting (Bio-Rad, Hercules, California, USA). The membranes were chemically cross-linked (Pall and Hamilton, 2008) and probed overnight with DNA or LNA (Locked Nucleic Acid probes biotinylated at the 5’ end (Supplementary Table 4). Detection was done using Chemiluminescent Nucleic Acid Detection Module (ThermoFischer, Waltman, MA, USA; Cat. Kit: 89880) and ChemiDoc equipment in Chemi High Resolution mode was used for image acquisition.

For the small RNA analysis, 10µg of total RNA was run on the 12% polyacrylamide gel as above, and the gel was stained with SYBR Gold. Signal was detected using the same equipment as above and image acquired using SYBR Gold mode.

### PCR

For RT-PCR, the extracted RNA was treated with DNAseI (Turbo DNAse, Thermoscientific) and purified again with the phenol/chloroform before cDNA synthesis using Revert Aid first strand cDNA synthesis kit (Thermoscientific) using oligo dT primer. qPCR was performed using Luminaris Color HiGreen qPCR Master Mix (Thermo Fisher Scientific) and performed on a LightCycler 96 Instrument (Roche) real-time PCR machine. Data were obtained from three independent biological replicates and were normalized to same references as described previously (Pratx et al., 2025).

### HTS sample preparation and data analysis

Caryopses ranging from 3 to 5 mm (3 to 5 DAF) were dissected from green developing spikes and the anthers have been removed (Supplementary Figure 3A). Three caryopses was bulked together to produce a sample; three samples were collected for each mutant line and 5 samples for wild type barley. Total RNAs were extracted from each sample same way as mentioned above for the northern blotting, divided into two and sent for high-throughput sequencing (HTS): small RNA-sequencing and polyA-selected RNA-sequencing in parallel. The raw sequencing data of HTS sequencing have been deposited in the SRA database (PRJNA1498662).

Matured green seeds of about 7 mm were collected from mother plants and endosperm isolated for total RNA extraction. Three endosperms were collected into one sample and five samples were collected both for the WT and *rdr2_2.* RNA samples were again divided into two and sent for sRNA sequencing and polyA-selected RNA sequencing. The raw sequencing data has been deposited in the SRA database.

Small RNA-sequencing reads were trimmed and filtered for quality with fastp bioinformatic tool (Langmead and Salzberg, 2012). The data quality of one *rdr2-1* caryopsis sample was not adequate, so it was excluded from the analysis. Further filtering was applied for tRNA, rRNA, snRNA and snoRNA specific reads using Bowtie2. 24-nt RNA clusters were identified and their expression data was calculated with ShortStack (Johnson et al., 2016).

PolyA-selected RNA-sequencing reads were trimmed and filtered for quality with Cutadapt (Kim et al., 2019). We have lost one sample of the *rdr2-3* caryopsis data due to low data quality. We used Hisat2 to align reads on barley genome, StringTie (Pertea et al., 2016) and Cuffdiff (Trapnell et al., 2013) for differential expression analysis. We created gene ontology plots with ShinyGO (v0.85.1; (Ge et al., 2020)).

We used MorexV3 genome assembly (version 62; (Yates et al., 2022)) available on the Ensembl Plants repository for both sequencing data analyses. PCAGO web tool (https://pcago.bioinf.uni-jena.de/) was used to create principal component analysis (PCA) plots from expression data.

### Analysis of grain morphology, composition and yield

The Marvin System (MarviTech GmbH, Wittenburg, Germany) was used for determination of the The Marvin System (MarviTech GmbH, Wittenburg, Germany) was used for determination of the thousand grain weight (TGW), grain width (GW) and grain length (GL) per individual plants according to the relevant industrial standard (MSZ 6367/4-86). For measuring the grain’s composition, 6-to-14 g of grain pools were milled using a ball mill (Retsch Mixer Mill MM 400), to produce wholemeal samples of control and mutant lines respectively. The total protein content of wholemeal samples was measured by the Dumas method with an Elementar Rapid N III Analyzer (Elementar Analysensysteme GmbH, Langenselbold, Germany) according to the ICC 167 standard method. The total amount of mixed-linkage β-glucan was determined in wholemeal samples using a Megazyme kit (Megazyme, Bray, Ireland) according to the AACC, Method 32–23.01 (1995). The amylose content of the starch was measured by Megazyme method (Yun and Matheson, 1990). Duplicate analyses were carried out on each sample and when the difference between them was higher than 10%, two more replications were measured.

Size-exlusion high-performance liquid chromatography (SE-HPLC) was used to determine the glutenin, gliadin, and albumin + globulin contents and the glutenin-to-gliadin ratio using a modification method (Batey et al., 1991), as described previously (Takac et al., 2021). Analysis has been carried out with three replicate extracts.

### HS treatment

Barley seeds were germinated in plastic box on wet filter paper. To investigate HS memory in barley seedlings were primed with a two-step acclimation (ACC; 38 for two hours-21 for one hour-44 for one hour) treatment 3 days before being exposed to a strong HS treatment (44 for three hours) on day 7 as described previously (Pratx et al., 2025). After HS the plantlets were moved into substrate pellet and further grown for twenty days at 21. At this stage the phenotypes of primed (ACC) and HS only WT and mutant plants were compared by measuring the length of the plants

## Data availability

Raw polyA RNA-seq and sRNA-seq data have been made available in SRA repository under the BioProject ID: PRJNA1498662.

## Acknowledgement

This work was supported by the Hungarian National Research, Development and Innovation Office (NKFIH), grant numbers ADVANCED 152238 (ZH), TKP2021-NKTA-06 (MR), NKFIH K137722 (TCS) and by the Flagship Research group Programme of the Hungarian University of Agriculture and Life Sciences. Tibor Csorba was supported by the Research Excellence Programme of the Hungarian University of Agriculture and Life Sciences. The biochemical analysis and research was carried out using the facilities of the Cereal Research Infrastructure Network (3-110-H).

## Author contributions

AA: genome editing, phenotyping, experimentation, data analyses, writing. ÁD: experimentation, writing, statistical analysis of phenotypic data. HMS: HTS data analyses. AK: genome editing. DP: phenotyping. MR: grain quality measurement. TC: writing, data analyses. ZH: conceptualization, supervision, writing, HS experiments.

## Supplementary Figures

**Supplementary Figure 1:**
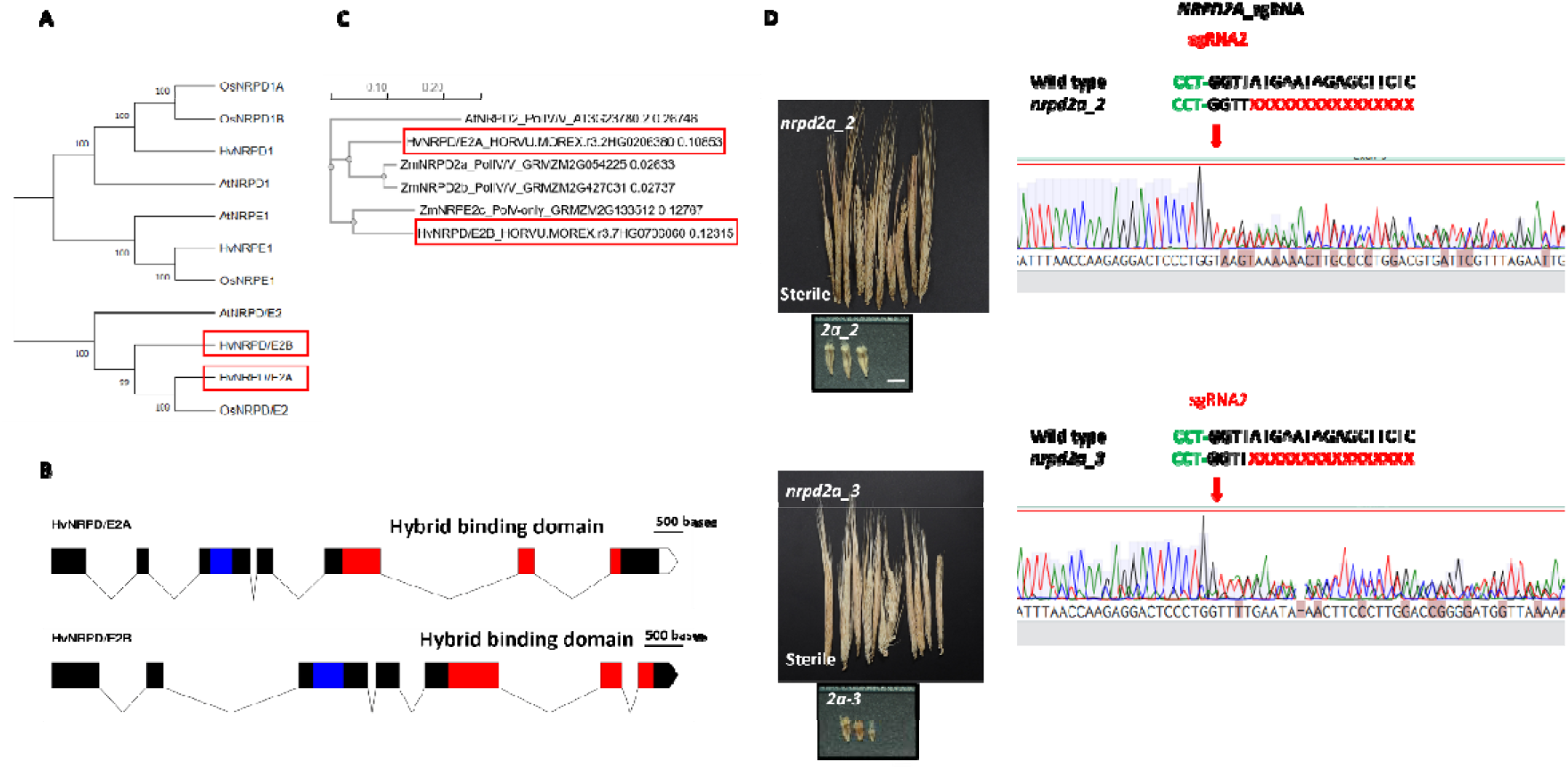
**A** Phylogenetic tree of *Arabidopsis*, rice and barley *NRPD* genes. **B** Schematic representation of barley *NRPD/E2* genes, solid black boxes represent exons and lines represents intron. Blue box shows the domain 2 also referred to as Lob domain. Red boxes show the hybrid binding domain also referred to as domain 6 which is the site of metal binding. The empty box shows 3’ UTR. **C** Phylogenetic tree of *Arabidopsis,* barley and maize and *NRPD/E2* genes. **D** Analyses of *nrpd/e2a_2* and of *nrpd/e2a_3* knock out lines exhibiting mixed mutations at sgRNA2 target sites and their phenotypes.

**Supplementary Figure 2:**
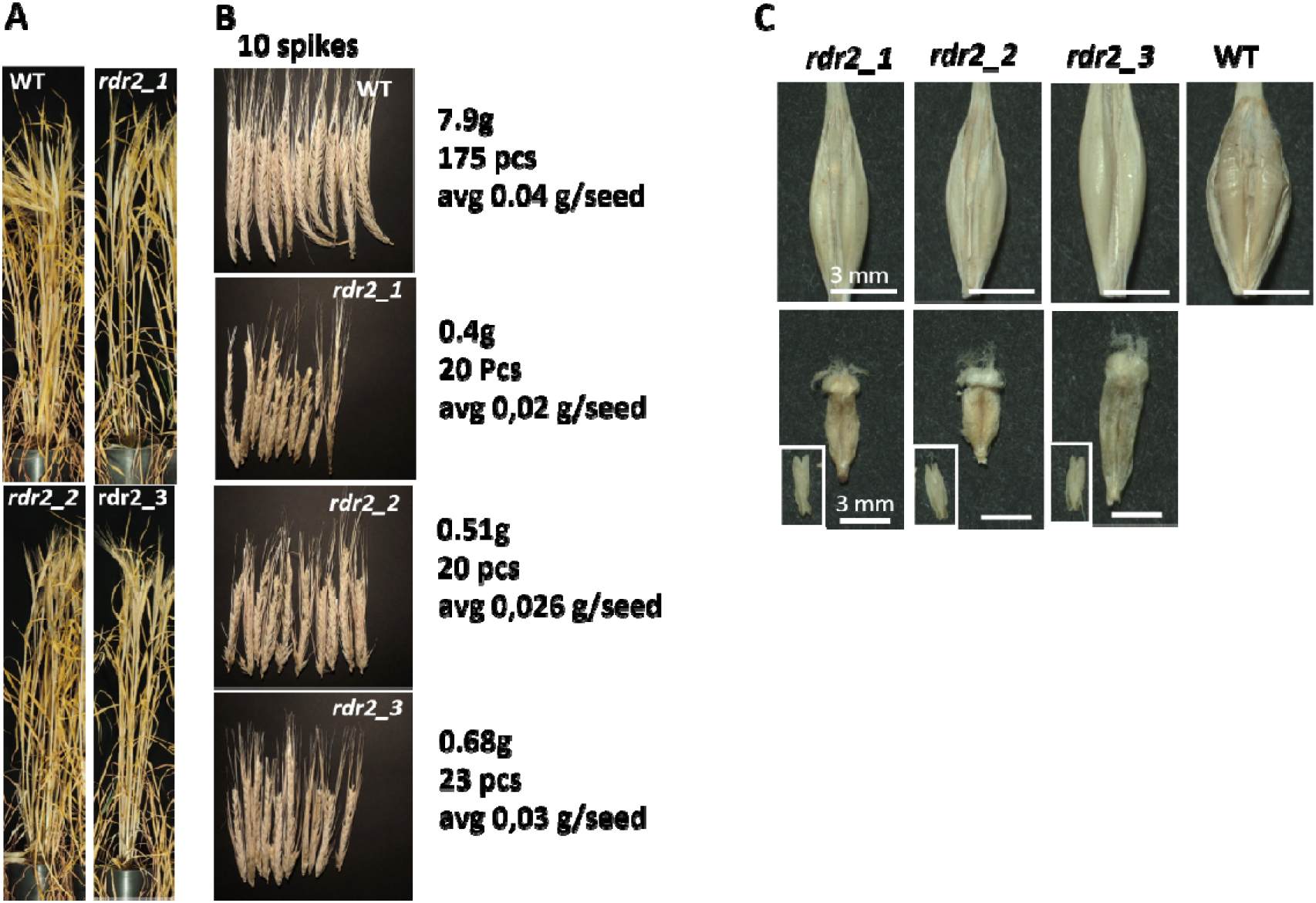
T1 generation of *rdr2* mutant lines exhibit seed development defects. **A** vegetative phenotype of mature dried wild type (WT) and *rdr2_1*, *rdr2_2* and *rdr2_3* plants. **B** 10 representative spikes of WT and mutant plants. The numbers on right show the weight of the seed production, the number of the seeds and average weight of the seeds. **C** The empty hulls of the mutant and the mature wild type seed (upper panels). The dissected aborted caryopses of the mutant lines. Insets show the developed anthers (lower panels).

**Supplementary Figure 3:**
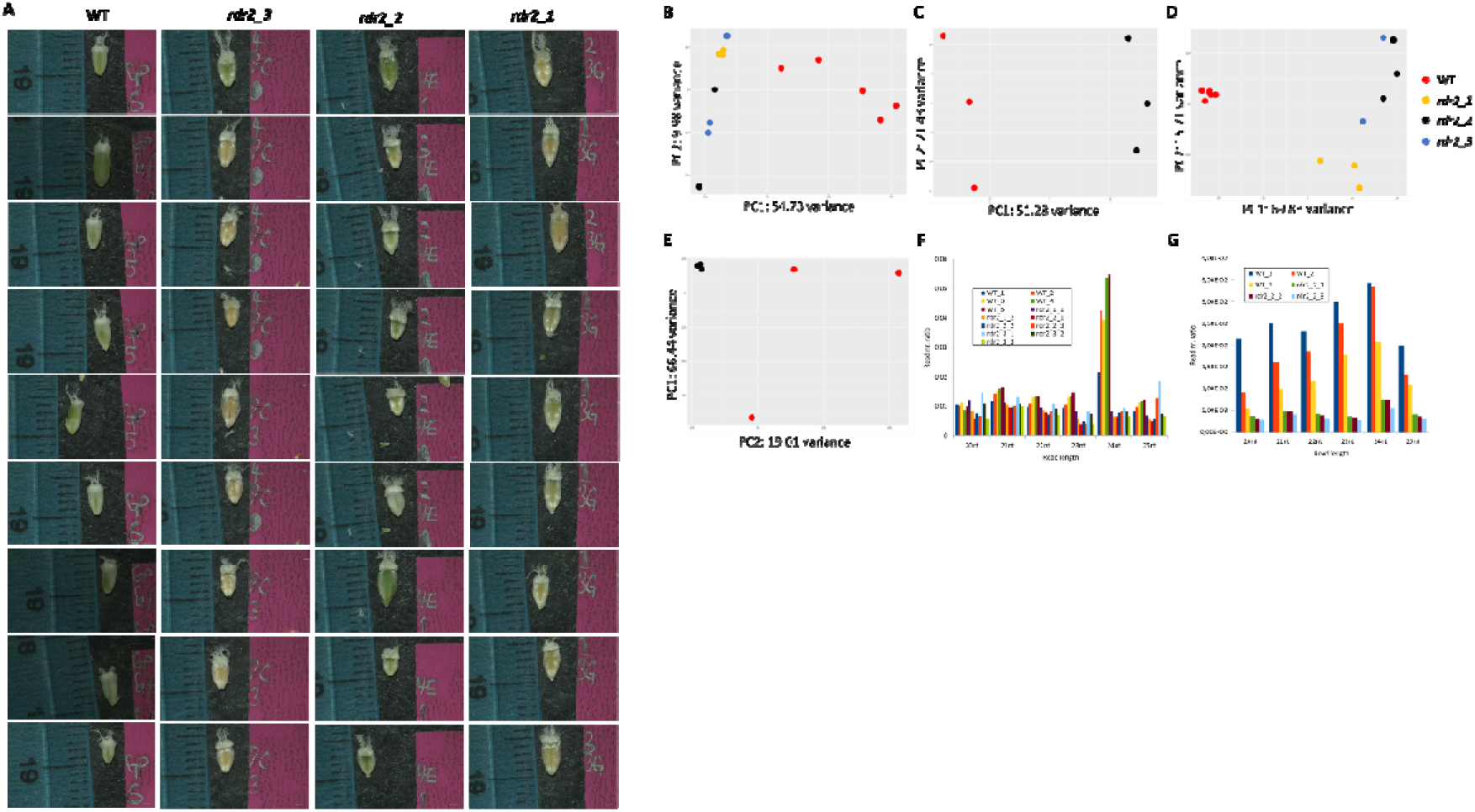
**A** Isolated caryopses for small RNA and mRNA HTS sequencing. Three caryopses were pooled in one sample. **B-E** Principal component analysis (PCA) from polyA-selected RNA sequencing of (**B**) caryopses and (**C**) endosperm, and from sRNA sequencing from (**D**) caryopses and (**E**) endosperm. **F-G** Ratio of reads with 20-25 nt of every sample in the small RNA sequencing data of (**F**) wild type (WT) and *rdr2-1*, *rdr2-2*, *rdr2-3* mutant caryopses or (**G**) wild type (WT) and *rdr2-1* mutant endosperm.

**Supplementary Figure 4:**
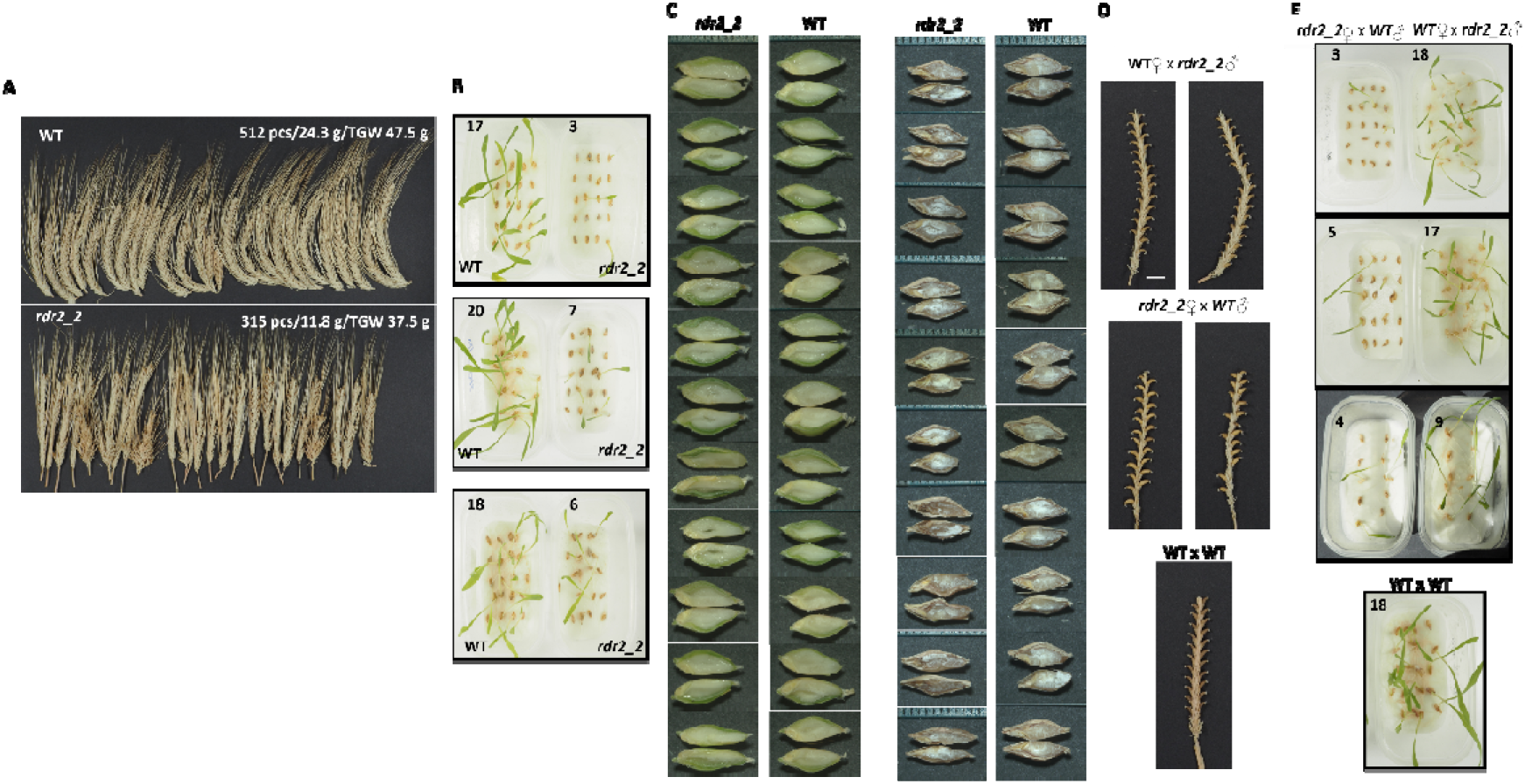
A. Another 25 representative spikes of WT and *rdr2_2* mutant, top right numbers show the number of seeds from the 25 spikes, weight of seeds and thousand grain weights respectively. **B** Germination efficiencies of WT *and rdr2_2* mutants in three replica experiments. **C** Dissected green seeds (left) and dried seeds (right) showing the abnormal endosperm in *rdr2_2.* **D** Dry spikes with seeds of replica experiments from the crossings of WT mother with *rdr2_2* pollen and *rdr2_2* mother with WT pollen. **D** Germination assays of seeds deriving from crossing experiments, in three replica experiments.

**Supplementary Figure 5.**
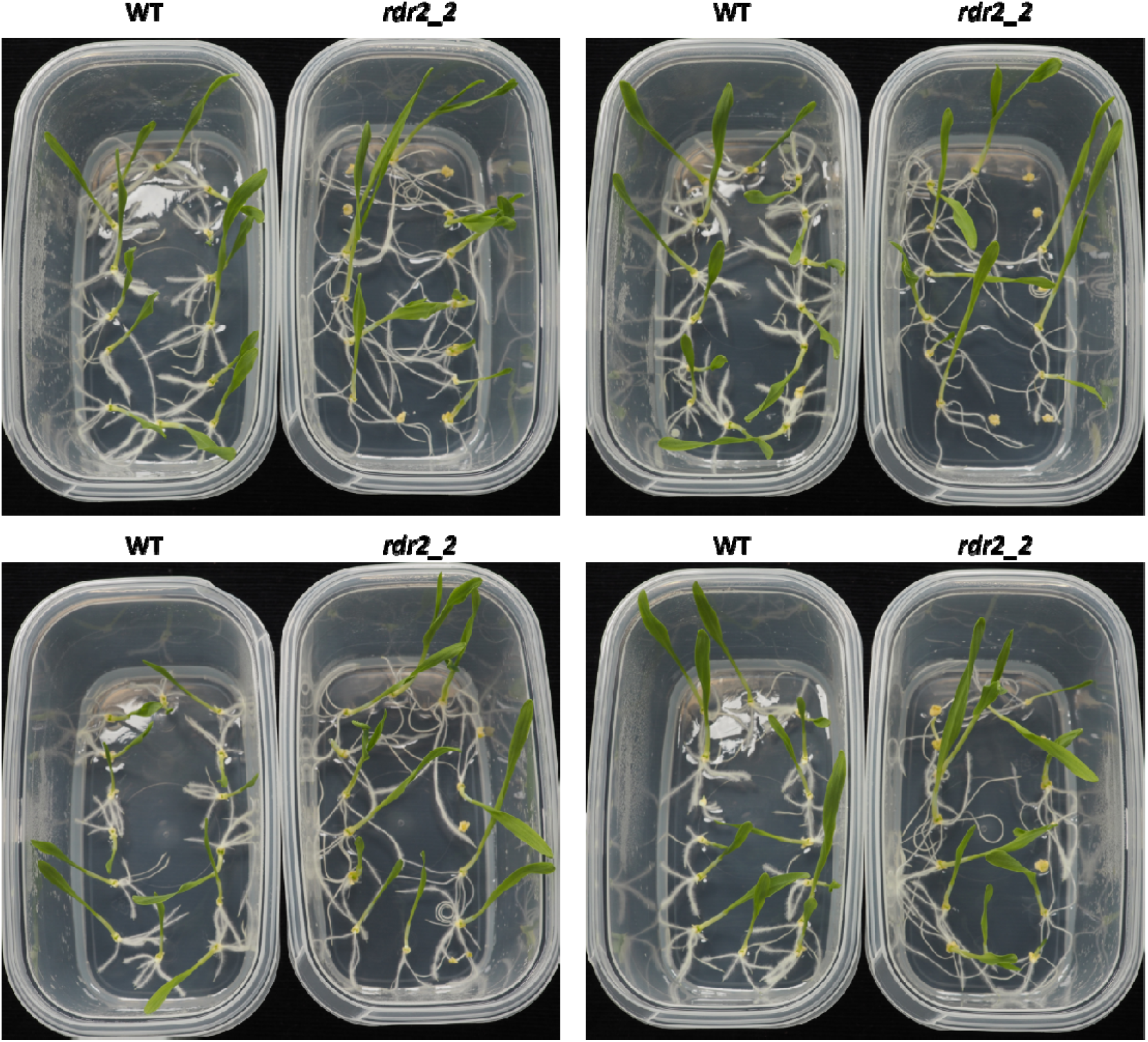
Embryo rescued wild type (WT) and mutant (*rdr2_2*) seeds germinated on medium. Replica experiments.

**Supplementary Figure 6:**
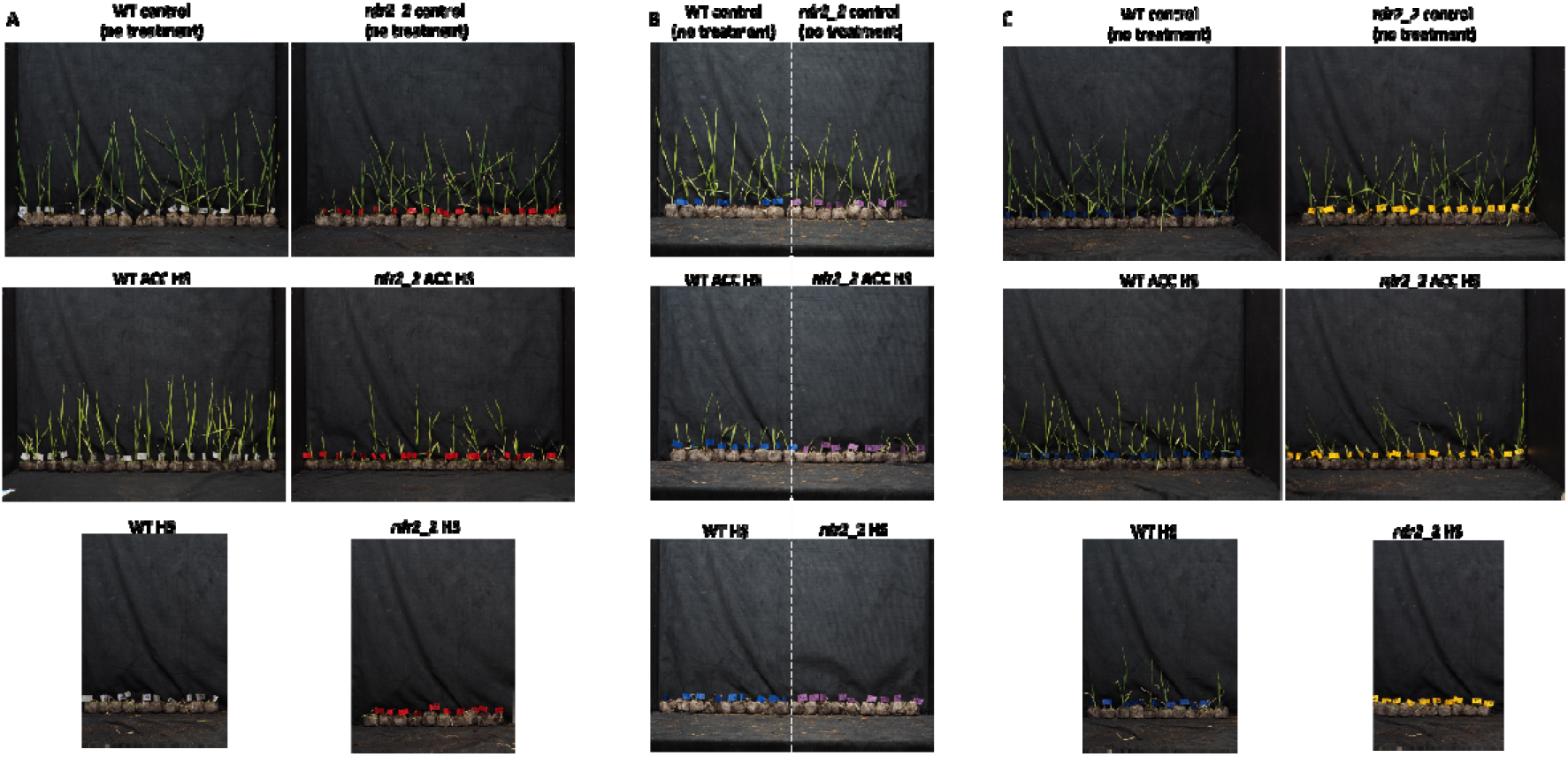
Representatives of replica heat shock (HS) experiments (A, B and C). Non treated control plants (upper panel). WT and *rdr2_2* plants received HS (44 for three hours) after three days of acclimation (ACC; 38 for two hours-21 for one hour-44 for one hour) (middle panel)). WT and *rdr2_2* plants received HS without ACC (HS) lower panel).

---

**Supplementary Table 2:** Physical and compositional traits of the wild type (WT) and mutant (*rdr2_2*) barley genotype (A) and their correlation to the kernel size (B).

|  | WT | rdr2_2 | WT SD | rdr2_2 SD | LSD 5% |
| --- | --- | --- | --- | --- | --- |
| TGW (g/1000kernel) | 47.57 | 35.57 | 0.2730 | 0.1097 | 0.6058 *** |
| Kernel width (mm) | 3.83 | 3.40 | 0.0577 | 0.0000 | 0.1755 ** |
| Kernel length (mm) | 8.43 | 8.87 | 0.0577 | 0.0577 | 0.1755 ** |
| Protein content % | 9.61 | 12.22 | 0.5111 | 0.0917 | 1.3284 ** |
| Glutelin/Prolamin ratio | 1.11 | 0.88 | 0.0395 | 0.0258 | 0.0431 ** |
| Albumin+Globulin/total % | 16.35 | 18.03 | 0.1545 | 0.9064 | 2.9424 n.s. |
| Prolamin/total % | 34.03 | 38.88 | 0.8307 | 0.2656 | 3.2251 * |
| Glutelin/total % | 37.64 | 34.05 | 0.8948 | 1.1114 | 3.1327 * |
| Amylose/amylopectin ratio | 0.40 | 0.38 | 0.0269 | 0.0171 | 0.1217 n.s. |
| Amylose content % | 28.42 | 27.76 | 1.3813 | 0.8953 | 6.2937 n.s. |
| Amylopectin content % | 71.58 | 72.24 | 1.3813 | 0.8953 | 6.2937 n.s. |
| β- glucan content (mg/g) | 43.52 | 45.53 | 1.1759 | 0.3473 | 2.7516 n.s. |

**B**
|  | TGW (g/1000kernel) |
| --- | --- |
| Protein content % | -0.9802 *** |
| Glutelin/Prolamin ratio | 0.9725 *** |
| Albumin+Globulin/total % | -0.8429 * |
| Prolamin/total % | -0.9765 *** |
| Glutelin/total % | 0.9108 ** |
| Amylose/amylopectin ratio | 0.3394 n.s. |
| Amylose content % | 0.3349 n.s. |
| Amylopectin content % | -0.3349 n.s. |
| β- glucan content (mg/g) | -0.8325 n.s. |

$r_{5\%}=0.7545$ , $r_{1\%}=0.8745$ , $r_{0.1\%}=0.9507$

